# Docking Molecular analysis of potential Aldosterone antagonists

**DOI:** 10.1101/2021.07.18.452814

**Authors:** Ivan Vito Ferrari, Paolo Patrizio

## Abstract

**Background:** Aldosterone antagonists (spironolactone, eplerenone) inhibit the action of aldosterone in the collecting duct; as such, these agents cause modest diuresis but inhibit potassium and hydrogen ion secretion. We report first time Potential Aldosterone antagonists by in Silico approach, using AutoDock Vina and AutoDock 4 (or MGL Tool), estimated with Pyrx and AM Dock Software, calculating three different important parameters: Binding Affinity (kcal/mol), estimated Ki (in nM units) and Ligand Efficiency (L.E. in kcal/mol). After a selective analysis of over 1000 drugs, processed with Pyrx (a Virtual Screening software for Computational Drug Discovery) in the Ligand Binding site pocket of the protein (ID PDB 2OAX Chain A:), we noticed high values of Binding Energy, about −13.55 kcal/mol estimated by AutoDock 4 with AM Dock Software, concluding that it could be an excellent candidate drug, compared to everyone else Aldosterone antagonists. Indeed, from the results of AutoDock Vina and AutoDock 4 (or AutoDock 4.2), implemented with Lamarckian genetic algorithm, LGA, trough AMDock Software, our results of Binding Energy are very similar to the crystallized Spironolactone in PDB 2OAX Chain A protein.

## 1. Introduction

Spirolactones are diuretics potent antagonists of the mineralocorticoid receptor (MR), a ligand-induced transcription factor belonging to the nuclear receptor superfamily. Spirolactones are synthetic molecules characterized by the presence of a C17 gamma-lactone, which is responsible for their antagonist character ^1^. hey may also be called aldosterone receptor blockers and they Aldosterone receptor antagonists may be used in the treatment of high blood pressure or heart failure. Aldosterone is the main mineralocorticoid hormone in the body and is produced in the adrenal cortex of the adrenal gland. Aldosterone increases sodium reabsorption by the kidneys, salivary glands, sweat glands and colon. At the same time, it increases the excretion of hydrogen and potassium ions. t plays a central role in the homeostatic regulation of blood pressure, plasma sodium (Na^+^), and potassium (K^+^) levels ^3–4^. By blocking the effects of aldosterone, aldosterone receptor antagonists block the reabsorption of sodium, which encourages water loss. Consequently, this leads to a decrease in blood pressure and a reduction in fluid around the heart ^4^. In this short communication, we investigated about 1000 drugs, through In Silico Docking approach, downloaded from PubChem Database (https://pubchem.ncbi.nlm.nih.gov/). In this paper particular attention, we have focused on AutoDock Vina, estimated with Pyrx software ^5^, a simple Virtual Screening library software for Computational Drug Discovery (https://pyrx.sourceforge.io/), based on prediction the binding orientation and affinity of a ligand and that of AutoDock ^6^, opened by AMDOCK software, (a versatile graphical tool for assisting molecular docking with AutoDock Vina and AutoDock 4, characterized to be automated docking of ligand to macromolecule by Lamarckian Genetic Algorithm and Empirical Free Energy Scoring Function ^7^.

## 2 Materials and methods

### 2.1 Protein and Ligand Preparation before docking

2OAX [Crystal structure of the S810L mutant mineralocorticoid receptor associated with SC9420] was prepared manually using different software, before molecular docking analysis. The first step, was downloaded from Protein Data Bank, https://www.rcsb.org/structure/2OAX and save in pdb format. The second step were removed all unnecessary docking chains. In fact, only the A chain (Mineralocorticoid receptor Chain A) has been maintained and re-saved in pdb format. **(See below figure 1)** Next, were the removal of ligands and water molecules crystallized using Chimera software ^8^. Later, polar hydrogens and Kohlmann charges were added with MGL Tool, (or called AutoDockTools, a software developed at the Molecular Graphics Laboratory (MGL) ^6–7^ As a last step they were added to the protein, any missing amino acids and the whole protein was minimized with the Swiss PDB Viewer Software ^9^. In this way the protein was saved in pdb format and ready for docking, through AutoDock Vina ^5^, estimated with a soft virtual screening library called Pyrx. On the other hand for Ligand Preparation, the first step, was to separate the crystallized drug from its protein, manually add all the hydrogens and their charges (with the MGL Tool software) and minimize it with MMFF94 force field ^10^, opened Pyrx software (https://pyrx.sourceforge.io/). This crystallized drug, Spironolactone was docked in the same binding area in order to accurately evaluate its Binding Affinity Vina Score with its protein and if there is overlapping of the drug Spironolactone, between its crystallized version and in its docking version. **(See below figure 2)** In the area of the drug crystallized with the protein’s A chain, more than 1000 dockings were carried out, through AutoDock Vina, estimated with Pyrx software, estimating their Binding Affinity (kcal/mol).

**Fig.1.**
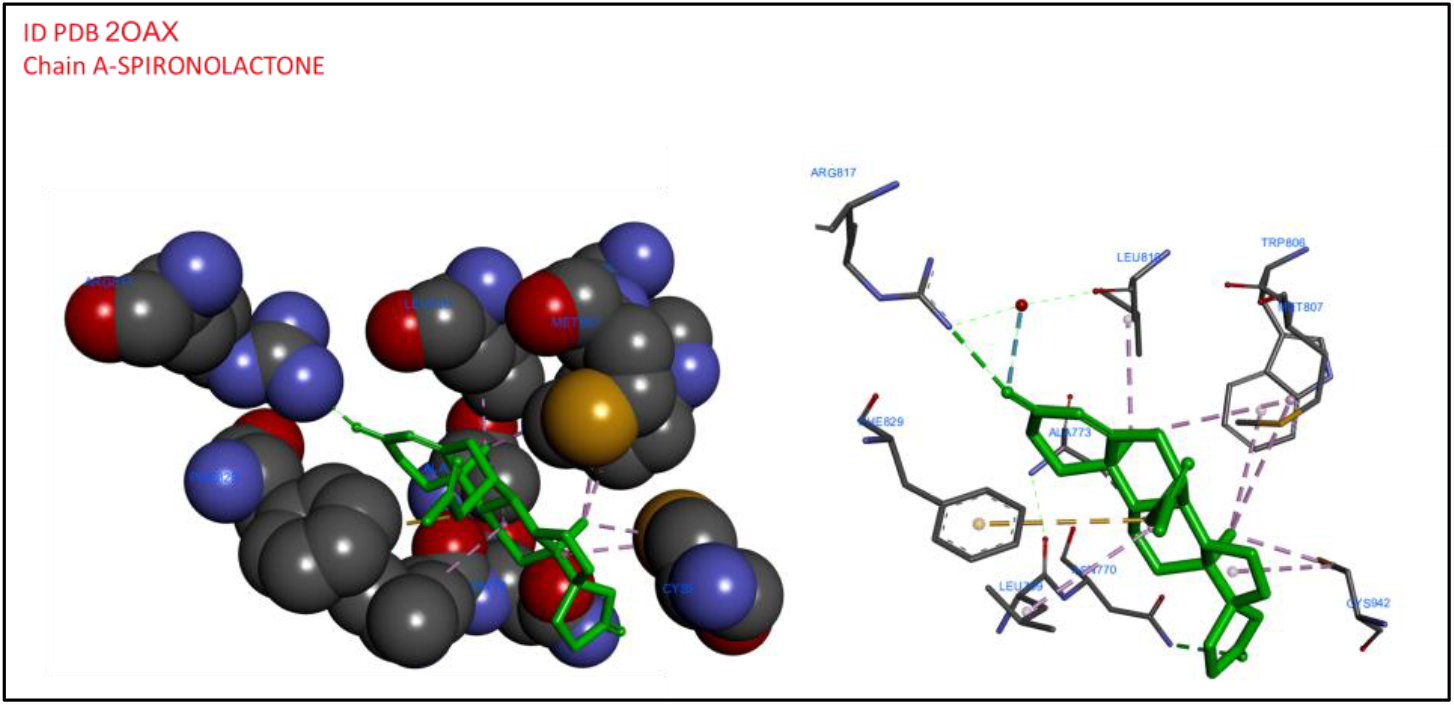
ID PDB 2OAX-complex Crystalized Spironolactone 3D Ligand Interactions characterized by Discovery Studio Biovia

**Fig. 2.**
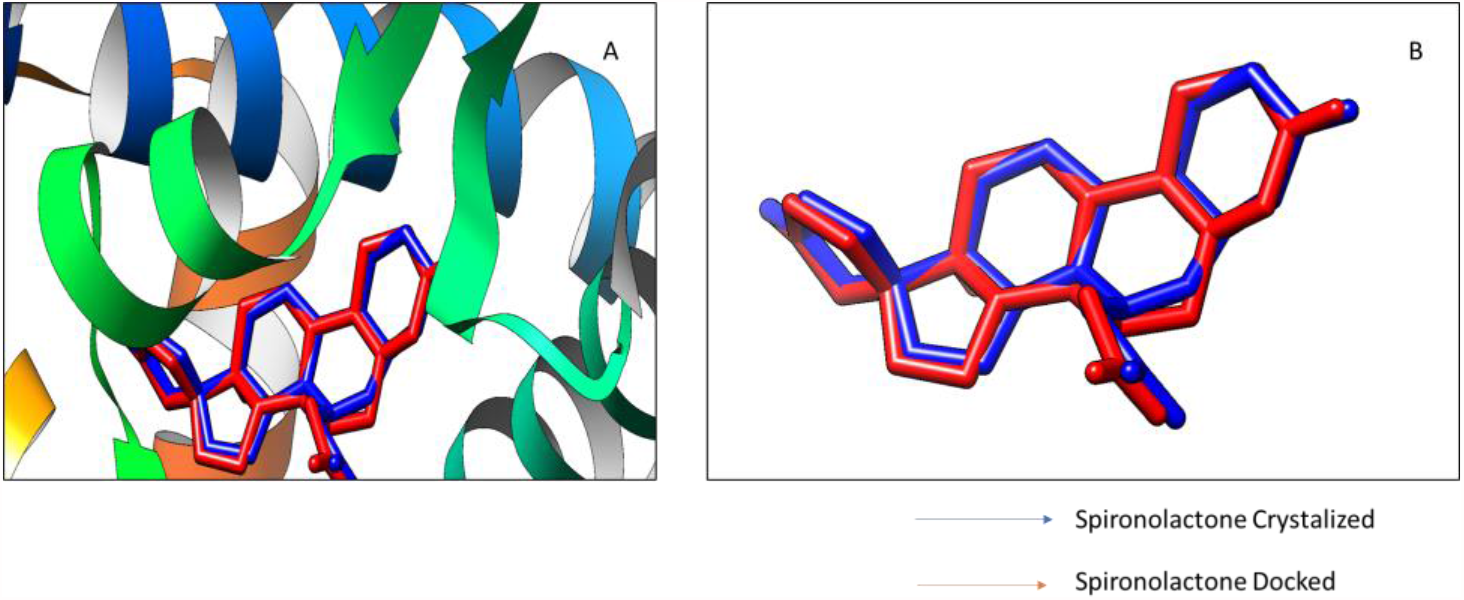
ID PDB 2OAX Comparison Crystalized Spironolactone with Spironolactone Docked in Ligand Binding Site Estimated by AutoDock Vina with Chimera Software

Parameters Grid Box for Docking in Ligand Binding Site Pocket for Repeatability Binding Affinity by AM Dock Software ^6^ calculated with AutoDock Vina and AutoDock 4:

- ID PDB ID PDB 2OAX Chain A: Center X (= −21.00); Centre Y (=60.00); Centre Z (=5.00); Dimensions (Angstrom) (Å) X, Y, Z [= 20=,20, =0]; exhaustiveness = 8

## 3 Discussion and Results

In this short communication we have investigated several natural compounds **(See below Table 1)** and drugs compounds, mainly potential Aldosterone antagonists,**(See below Table 2)**, by docking analysis approach in PDB 2OAX, studying their relative binding energies compared to crystallized Spironolactone. From our results we have shown, as expected, high Binding Affinity values kcal/mol of the Aldosterone antagonists, especially Mespirenone, 15β,16β-methylenespironolactone, is a steroidal antimineralocorticoid of the spirolactone group related to spironolactone that was never marketed ^11^. After a selective analysis of over 500 drugs, processed with Pyrx (a Virtual Screening software for Computational Drug Discovery) in the Ligand Binding site pocket of the protein (ID PDB 2OAX chain A), we noticed high values of these 3 parameters mentioned above of Mespirenone, concluding that it could be an excellent candidate drug used for potential Aldosterone antagonist. Indeed, from the results of AutoDock Vina and AutoDock 4 (or AutoDock 4.2), implemented with Lamarckian genetic algorithm, LGA, trough AM Dock Software. This a steroidal antimineralocorticoid both by AutoDock Vina and AutoDock 4 has excellent Binding affinity value, ca. −13,00 kcal/mol, a Ki value 0.21 nM and Ligand efficiency ca −0.45 kcal/mol. These results are comparable to the drug crystallized in the Spironolactone protein. **(See below figure 3–4)**

**Tab 1.** ID PDB 2OAX : “Crystal structure of the S810L mutant mineralocorticoid receptor associated with SC9420” of Best Natural Compounds Docking Binding Energy Score (kcal/mol) estimated by AutoDock Vina with Pyrx Software

| Ligand | Binding Energy (kcal/mol) |
| --- | --- |
| (-)-Epicatechin_uff_E=230.82 | -8.3 |
| 1-O-Acetylbritannilactone_uff_E=433.92 | -7.7 |
| Allicin_uff_E=116.06 | -4.4 |
| Amygdalin_uff_E=518.90 | -6.6 |
| Apigenin_uff_E=234.36 | -8.8 |
| Astragalin_uff_E=604.21 | -6.7 |
| Baicalein_uff_E=241.52 | -8.6 |
| Bilobalide_uff_E=817.34 | -6.7 |
| Borneol_uff_E=466.93 | -6.3 |
| Caffeine_uff_E=351.11 | -6.2 |
| Equol_uff_E=174.65 | -8.7 |
| Eucalyptol_uff_E=344.57 | -6.0 |
| Eupatorin_uff_E=492.88 | -6.7 |
| Fisetin_uff_E=344.71 | **-9.0** |
| Genistein_uff_E=356.74 | **-8.7** |
| Gingerol_uff_E=166.58 | -7.3 |
| Glabridin_uff_E=227.93 | -8.1 |
| Glycyrrhizin_uff_E=1232.94 | 21.3 |
| Hesperidin_uff_E=588.84 | 3.3 |
| Hexahydrothymol_uff_E=94.75 | -5.7 |
| Kaempferol_uff_E=362.50 | -8.3 |
| Limonene_uff_E=102.80 | -5.7 |
| Luteolin_uff_E=242.10 | **-8.9** |
| Magnolol_uff_E=333.60 | -7.5 |
| Myrcene_uff_E=90.40 | -5.4 |
| naringenin_uff_E=195.80 | -8.7 |
| Orcinol_uff_E=82.15 | -5.6 |
| Paraxanthine_uff_E=335.71 | -5.9 |
| Piceatannol_uff_E=172.68 | -8.1 |
| Quercetin_uff_E=379.98 | **-8.9** |
| Resveratrol_uff_E=172.23 | -8.0 |
| Rutin_uff_E=754.29 | -3.7 |
| Salicin_uff_E=324.41 | -7.3 |
| Shogaol_uff_E=155.45 | -7.0 |
| Silibinin_uff_E=443.52 | 8.0 |
| Squalene_uff_E=256.38 | -7.3 |
| Theobromine_uff_E=330.95 | -5.8 |
| Theophylline_uff_E=338.65 | -6.0 |
| Zingiberene_uff_E=205.48 | -6.8 |

**Tab 2.** ID PDB 2OAX : “Crystal structure of the S810L mutant mineralocorticoid receptor associated with SC9420”, of Best Drugs Docking Binding Energy Score (kcal/mol) estimated by AutoDock Vina with Pyrx Software

| Ligand | Binding Energy (Kcal/mol) |
| --- | --- |
| Abiraterone_mmff94_E=109.51 | -9.9 |
| Prednisolone_mmff94_E=133.96 | -9.6 |
| Prednisone_mmff94_E=121.67 | -9.6 |
| Alfatradiol_mmff94_E=59.38 | -9.8 |
| Sitagliptin_mmff94_E=41.81 | -9.9 |
| Betamethasone_mmff94_E=187.30 | -10.0 |
| Valsartan_mmff94_E=69.88 | -9.6 |
| Vamorolone_mmff94_E=132.02 | -9.7 |
| Clobetasol_mmff94_E=174.79 | -10.3 |
| SPIRONOLACTONE CRYSTALIZED mmff94_E=429.27 | -12.5 |
| Cortisone_mmff94_E=119.79 | -9.9 |
| Cortodoxone_mmff94_E=124.34 | -10.3 |
| Hydrocortisone_mmff94_E=130.65 | -9.8 |
| Hydroxyprogesterone_mmff94_E=102.34 | -10.4 |
| Dehydrocorticosterone_mmff94_E=94.26 | -10.5 |
| Desoximetasone_mmff94_E=151.31 | -9.8 |
| Methylprednisolone_mmff94_E=142.36 | -9.6 |
| Methyltestosterone_mmff94_E=81.23 | -10.3 |
| Medroxyprogesterone_mmff94_E=107.33 | -10.7 |
| Mespirenone_mmff94_E=103.34 | -13.1 |
| Duraspiron_mmff94_E=79.94 | -10.2 |
| Estradiol_mmff94_E=58.52 | -10.0 |
| Ethinyl_estradiol_mmff94_E=61.61 | -10.4 |
| Fludrocortisone_mmff94_E=156.61 | -10.4 |
| Fluoroestradiol_F-18_mmff94_E=77.18 | -10.2 |
| Fluoxymesterone_mmff94_E=117.33 | -10.3 |

**Fig. 3.**
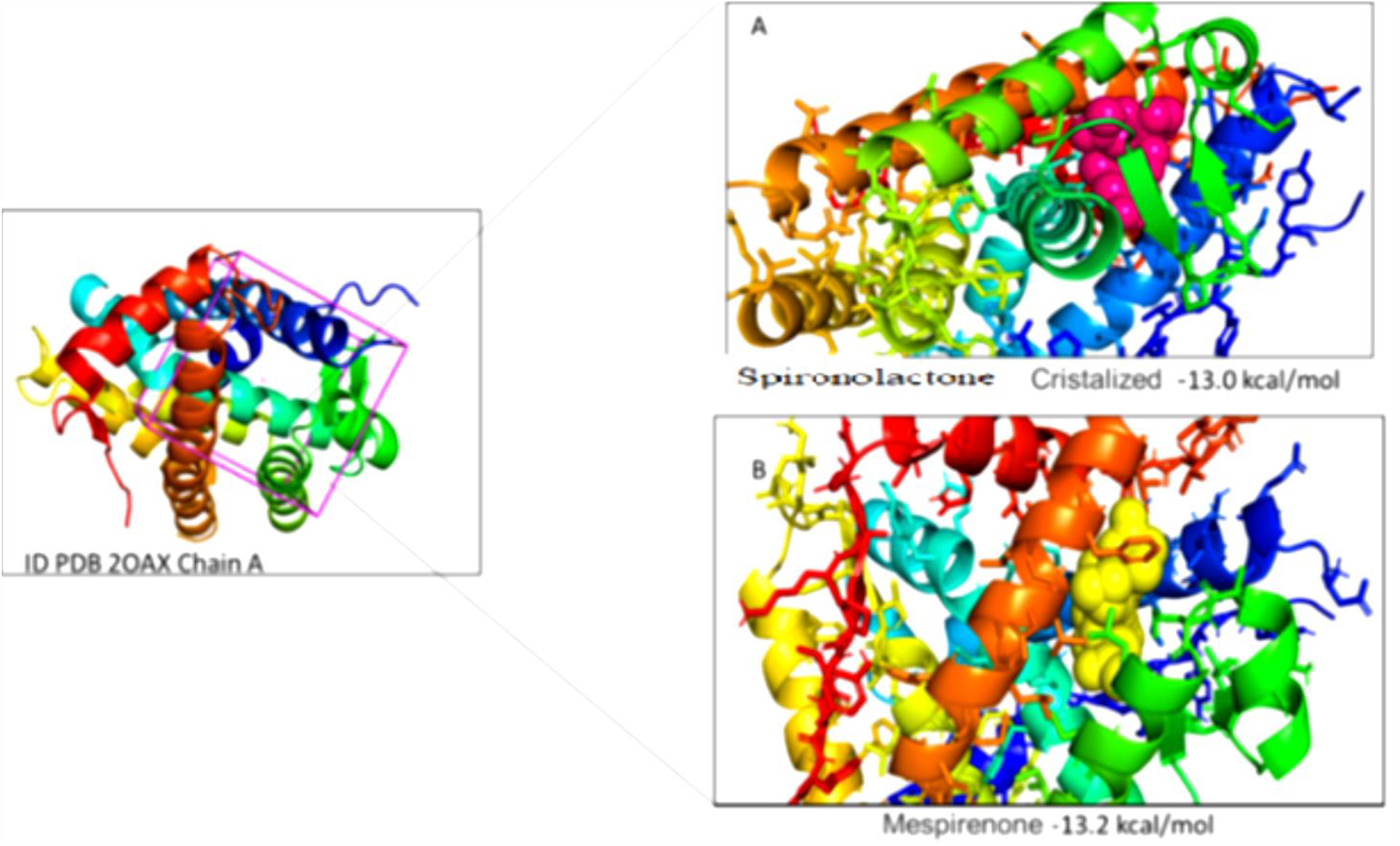
Docking Results ID PDB 2OAX Chain A: a) Purple sphere Spironolactone Crystalized −13.00 kcal/mol b) yellow sphere Mespirenone −13.2 kcal/mol estimated by AutoDock Vina with AM Dock Software

**Fig. 4.**
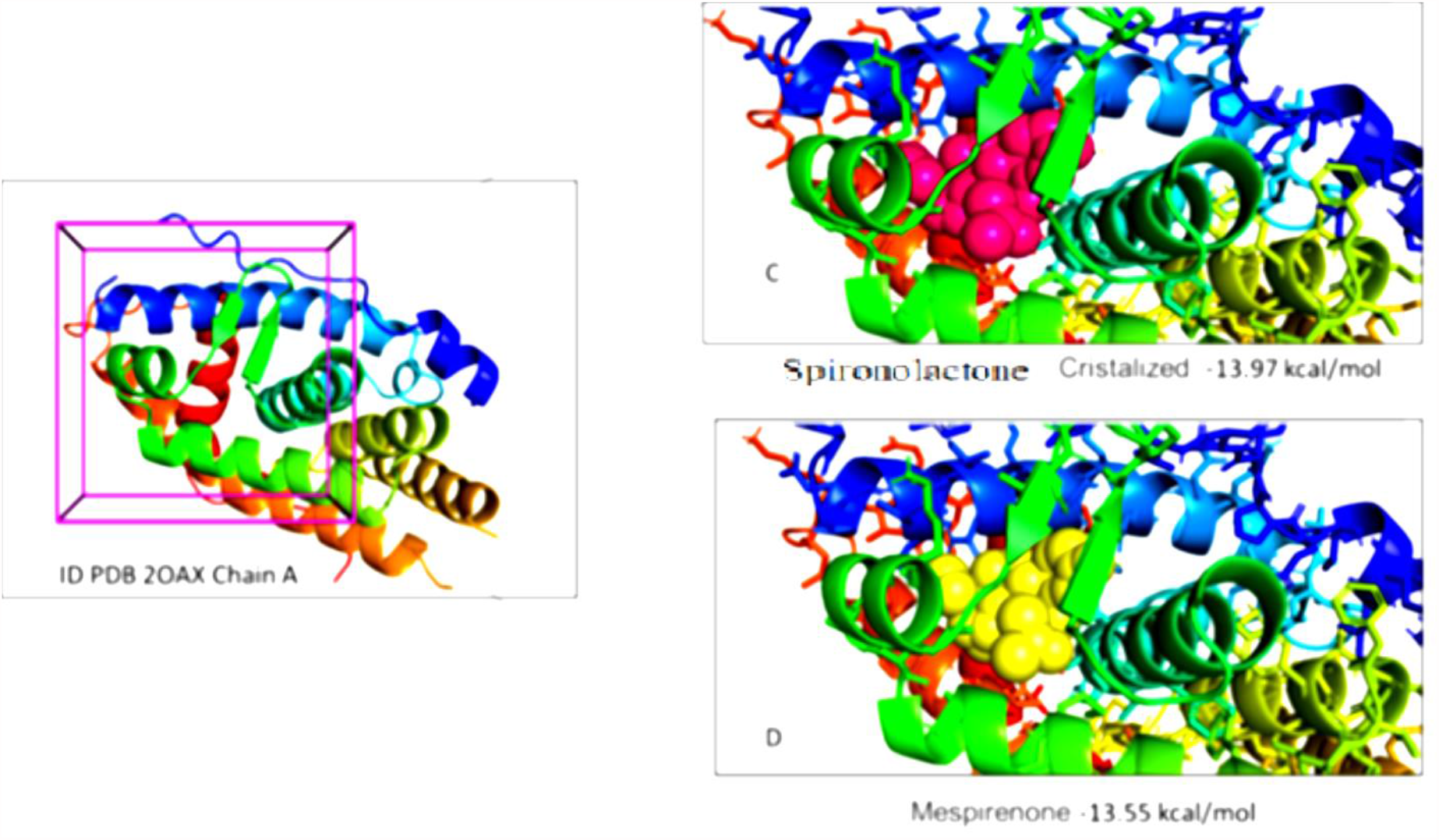
Docking Results ID PDB 2OAX Chain A: c) Purple sphere Spironolactone Crystalized −13.97 kcal/mol d) Yellow sphere Mespirenone −13.55 kcal/mol estimated by AutoDock 4 with AM Dock Software

## 4. Conclusion

In this short communication we have focused on the role of Aldosterone antagonists, mainly we have investigated by in Silico approach a potential capacity of Mespirenone. It is a spironolactone derivative and aldosterone antagonist with diuretic activity. From our results of docking analysis are both Mespirenone and Spironolactone are comparable Binding Energy in Ligand Binding Site pocket of protein ID PDB 2OAX, about −13.00 kcal/mol. From these preliminary computational results, future in vitro and in vitro research will be possible, to understand better their effective function of Aldosterone antagonists.

## Conflict of interest declarations

## Notes

### Competing Interest Statement

The authors have declared no competing interest.

## References

1. Huyet, J., Pinon, G. M., Fay, M. R., Fagart, J., & Rafestin-Oblin, M. E. (2007). Structural basis of spirolactone recognition by the mineralocorticoid receptor. Molecular pharmacology, 72(3), 563–571.

2. Jaisser, F., & Farman, N. (2016). Emerging roles of the mineralocorticoid receptor in pathology: toward new paradigms in clinical

3. Marieb, E. N., & Hoehn, K. (2007). Human anatomy & physiology. Pearson education.

4. https://www.drugs.com/drug-class/aldosterone-receptor-antagonists.html

5. Trott, Oleg, and Arthur J. Olson. “AutoDock Vina: improving the speed and accuracy of docking with a new scoring function, efficient optimization, and multithreading.” Journal of computational chemistry 31.2 (2010): 455–461

6. Morris, G. M., Huey, R., Lindstrom, W., Sanner, M. F., Belew, R. K., Goodsell, D. S. and Olson, A. J. (2009) Autodock4 and AutoDockTools4: automated docking with selective receptor flexiblity. J. Computational Chemistry 2009, 16: 2785–91.

7. http://mgltools.scripps.edu/

8. Pettersen, Eric F., et al. “UCSF Chimera—a visualization system for exploratory research and analysis.” Journal of computational chemistry 25.13 (2004): 1605–1612.

9. Guex, Nicolas, and Manuel C. Peitsch. “SWISS-MODEL and the Swiss-Pdb Viewer: an environment for comparative protein modeling.” electrophoresis 18.15 (1997): 2714–2723.

10. Thomas A. Halgren, J. Comput. Chem., 17, 490–519 (1996).

11. Losert, W., Bittler, D., Buse, M., Casals-Stenzel, J., Haberey, M., Laurent, H., … & Wiechert, R. (1986). Mespirenone and other 15, 16-methylene-17-spirolactones, a new type of steroidal aldosterone antagonists. Arzneimittel-forschung, 36(11), 1583–1600.

